# Altered theta / beta frequency synchrony links abnormal anxiety-related behavior to synaptic inhibition in Neuroligin-2 knockout mice

**DOI:** 10.1101/726190

**Authors:** Hugo Cruces-Solis, Olga Babaev, Heba Ali, Carolina Piletti Chatain, Vasyl Mykytiuk, Nursen Balekoglu, Sally Wenger, Dilja Krueger-Burg

## Abstract

Inhibitory synaptic transmission plays a key role in the circuits underlying anxiety behaviors, but the network mechanisms by which disruptions in synaptic inhibition contribute to pathological anxiety processing remain largely unknown. Here we addressed this question in mice lacking the inhibitory synapse-specific adhesion protein Neuroligin-2 (Nlgn2), which display widespread reduction in inhibitory synaptic transmission as well as a pronounced anxiety phenotype. To investigate how the lack of synaptic inhibition alters the communication between key brain regions in anxiety processing, we recorded local field potentials (LFPs) simultaneously from a network of brain regions involved in anxiety processing, including the basolateral amygdala (BLA), centromedial amygdala, bed nucleus of the stria terminalis, prefrontal cortex and ventral hippocampus (vHPC). We found that LFP power in the vHPC was profoundly increased while vHPC-directed theta frequency synchrony was disrupted in Nlgn2 KO mice under anxiogenic conditions. Instead, deletion of Nlgn2 increased beta frequency synchrony across the anxiety network, and the theta / beta synchrony ratio strongly predicted anxiety behaviors in an open field paradigm. Local deletion of Nlgn2 in the vHPC and BLA revealed that they encode distinct aspects of the anxiety phenotype of the Nlgn2 KO mice, with vHPC linked to anxiety induced freezing and BLA linked to reduction in exploratory activity. Together, our data demonstrate that alterations in long-range functional connectivity link synaptic inhibition to abnormal anxiety behaviors, and that Nlgn2 KO mice represent an interesting model to study the role of inhibitory synaptic transmission in the circuits underlying anxiety disorders.

## Introduction

Anxiety disorders are among the most common psychiatric illnesses, affecting an estimated one in nine people in any given year and posing a significant burden for affected individuals, their families and society (1, 2). Pathological anxiety arises when normally adaptive defensive behaviors become persistent, excessive and out of proportion to the actual threat posed, severely interfering with activities of daily life (1–5). Understanding the biological mechanisms that underlie the transition from adaptive to pathological anxiety is therefore paramount in developing new treatment strategies for anxiety disorders in order to alleviate this burden.

One of the factors that contribute to the development of pathological anxiety behaviors are alterations in GABAergic inhibitory synaptic transmission (6, 7). Anxiolytic medications such as benzodiazepines target the GABAergic system, and variants in components of the molecular machinery at inhibitory synapses have been associated with anxiety disorders (5–8). One of these components is Neuroligin-2 (Nlgn2), a postsynaptic adhesion protein that is localized selectively to inhibitory synapses (8–10) and that has been linked to anxiety disorders and co-morbid autism and schizophrenia (11, 12). Deletion of Nlgn2 in mice results in impaired inhibitory synaptic transmission in the hippocampus, amygdala and cortex (8, 10, 13–16) as well as a profoundly abnormal anxiety phenotype (15, 17, 18). These observations indicate that loss of inhibitory synaptic transmission in the Nlgn2 knockout (KO) mice results in aberrant activation of the anxiety network, providing a unique opportunity to study the abnormal processing of anxiety information across the brain. In particular, we sought to determine whether these pathological anxiety-related behaviors are encoded in the same way as adaptive anxiety and simply represent an excessive activation of the relevant circuits, or whether the etiological factors that result in anxiety disorders may fundamentally alter information processing in these circuits.

To address this question, we performed simultaneous local field potential (LFP) recordings in multiple brain regions in wildtype (WT) and Nlgn2 KO mice under anxiogenic conditions in an open field (OF) paradigm. Specifically, we focused on five brain regions known to be essential for anxiety processing and the context-dependent regulation of anxiety-related behaviors, including the basolateral amygdala (BLA), centromedial amygdala (CeM), ventral hippocampus (vHPC), medial prefrontal cortex (mPFC) and bed nucleus of stria terminalis (BNST) (4, 19–27). Our data indicate that disruptions in synaptic inhibition due to loss of Nlgn2 result in a fundamental change in the mechanisms by which anxiety-related behaviors are encoded in the anxiety network.

## Materials and methods

### Animals

Nlgn2 KO mice (16) were generated in our laboratory on an 129/Sv background and were backcrossed onto a C57BL/6JRj background (Janvier Labs) for at least six generations. For experiments, male WT and KO littermates were generated from heterozygous breeders. Conditional Nlgn2 KO mice were generated in our laboratory on a C57BL/6N background (see Figure S1 and S2 for details of generation and validation) and were backcrossed onto a C57BL/6JRj background for at least six generations. For experiments, male homozygous cKO mice were generated from homozygous breeders. Mice were 8-12 weeks old at the beginning of the experiment. All animals were maintained on a 12 hour light/dark cycle (7 am / 7 pm), with food and water *ad libitum*, and all experiments were performed during the light cycle. The experimenter was blind to genotype during all stages of data acquisition and analysis. All procedures were approved by the State of Niedersachsen (Landesamt für Verbraucherschutz und Lebensmittelsicherheit) and were carried out in agreement with the guidelines for the welfare of experimental animals issued by the Federal Government of Germany and the Max Planck Society.

### Electrode implantation for *in vivo* electrophysiology

Mice received an intraperitoneal (i.p.) injection of Carprofen (5 mg/kg) to reduce post-surgery pain 30 min prior to surgery, and they were then anesthetized with avertin (loading dose 2 mL/100 g, maintenance doze 0.2 mL/100 g intraperitoneal). Anesthetized mice were placed in a digital stereotaxic frame and their body temperature was monitored by a rectal probe and maintained at 36 °C. An incision in the midline of the scalp was made to expose the skull. Bregma and lambda were aligned to a plane level ± 50 µm. Electrodes were implanted individually into the right vHPC (−3.08 mm AP, 3.4 mm ML and 3.8 mm DV), mPFC (2.1 mm AP, 0.4 mm ML and 1.4 DV), BLA (−1.22 mm AP, 3.15 mm ML and 4.3 mm DV), CeM (−1.22 mm AP, 2.3 mm ML and −5.04 mm DV) and BNST (−0.2 mm AP, 0.4 mm ML and 4.3 mm DV). The electrodes consisted of individual insulated tungsten wires (70 µm inner diameter) inserted into a polymide tube (127 µm inner diameter) and attached to an 18-pin Omnetics connector. A reference screw was implanted above the cerebellum. The implant was secured with three screws implanted in the skull and bonded with dental cement.

Immediately after the surgery, mice received subcutaneously analgesic (Carprofen 5 mg/kg) and antibiotic (Baytril 5 mg/kg). 24 hours after the surgery, mice received Carprofen (5 mg/kg) subcutaneously and Baytril in the drinking water (0.2 mg/ml). After surgery, mice were single-housed and were allowed to recover for 7 days before behavioral analysis and data acquisition (see below). They were then sacrificed for histological verification of the recording site (Figure S3).

### *In vivo* electrophysiology data acquisition during OF behavior

OF behavior and *in vivo* electrophysiological recordings were performed after 7 days of recovery. Mice were first habituated to the recording setup for four days, during which they were tethered to the acquisition system and recorded for five minutes in a habituation context (home cage with dim light ~ 30 lux). On the fifth day, mice were tethered to the system and recorded during five minutes in the habituation condition. Afterwards, the mice were placed into a wooden circular open field chamber (50 cm diameter) with a light intensity of 90 lux, and they were permitted to explore the chamber for 15 min. Between each mouse, the arena was cleaned thoroughly with 70% ethanol followed by water to eliminate any odors left by the previous mouse.

Electrophysiological signals were amplified, digitized, multiplexed and sent to the acquisition board. We used a RHD2132 16-channel amplifier board (Intan Technologies) connected to the Open Ephys acquisition system via an ultra-thin SPI interface cable (Intan Technologies). The raw signal was acquired at 30 kHz sampling rate, bandpass filtered (0.1-7500 Hz) and stored for offline analysis. During the experiment, simultaneous electrophysiological and video recordings were acquired with the Open Ephys GUI software.

### Data analysis for *in vivo* electrophysiology

The Bonsai software (28) was used to extract the location of the mouse in each video frame. The set of coordinates were imported to MATLAB, where location and speed were calculated. Local field potentials (LFPs) were analyzed in custom-written MATLAB scripts. The signal was filtered between 0.7 and 400 Hz using a zero-phase distortion FIR filter and down sampled to 1 kHz. To exclude potential locomotion influences in the LFP recordings, we only analyzed periods of exploration with a speed > 7 cm/s. To this end, we extracted the electrophysiological traces in which the animals were moving with a speed of 7 to 40 cm/s in the habituation condition and periphery of the open field and concatenated them into one continuous trace. The power spectrum was calculated using the *pwelch* function in MATLAB with 1 s windows and 0.8 s overlap. For coherence we computed the imaginary part of the cross power spectral density normalized by the square root of the product of the auto power spectral densities, using the same window and overlap. For each frequency band, we used the following frequency ranges: theta: 6-12 Hz; beta: 15-30 Hz; gamma: 35-70 Hz. To obtain an individual value of power for each animal, we summed the power within a given frequency band, while for coherence we computed the maximum value within a frequency band. For the power and coherence correlations as a function of time spent in the center, we used the power and coherence values computed for each mouse and the linear correlation coefficient (*fitlm* function, MATLAB) was calculated including mice from all genotypes. The Granger causality was calculated using the Multivariate Granger Causality Toolbox (29, 30). We used the same unfiltered LFP traces used for coherence and power calculations. The order of the vector autoregressive (VAR) model to be fitted was calculated using the Akaike information criterion.

### Generation of AAV-GFP and AAV-GFP/Cre viruses

To locally delete Nlgn2 using local expression of Cre recombinase, we generated AAV vectors using the AAV-GFP and AAV-GFP/Cre plasmids generously provided by Fred Gage (31) (Addgene plasmids 49055 and 49056) in combination with the pDP5rs packaging system (PlasmidFactory, Bielefeld, Germany) as previously described (18). Briefly, plasmids were transfected into HEK cells using polyethylenimine transfection, and virus particles were harvested 48-72 hours later. To this end, HEK cells were lysed for 30 min at 37°C in 20 mM Tris, pH 8.0, containing 150 mM NaCl, 0.5% sodium deoxycholate and benzonase, followed by incubation in 1 M NaCl at 56°C for 30 min. Lysates were stored at −80°C overnight, thawed, and purified on a 15%/25%/40%/54% iodixanol gradient by ultracentrifugation (90 min at 370,000 x g). The 40% fraction was collected, diluted in PBS containing 1 mM MgCl_2_ and 2.5 mM KCl, and concentrated using an Amicon 100K MWCO filter.

### Stereotaxic injection of AAV-GFP and AAV-GFP/Cre viruses

Mice were anesthetized with avertin (loading dose 2 mL/100 g, maintenance doze 0.2 mL/100 g intraperitoneal). Anesthetized mice were placed in a digital stereotaxic frame, and an incision in the midline of the scalp was made to expose the skull. Bregma and lambda were aligned to a plane level ± 50 µm. 0.8 μl of virus was injected bilaterally into the BLA (−0.8 mm AP, ±3.2 mm ML and −5.2 mm DV) or the vHPC (−2.3 mm AP, ±3.2 mm ML and −4.2 mm DV) using a Nanoject II Microinjector (Drummond, Broomall, PA, USA) and a Micro pump controller (WPI). Mice were alternately assigned to receive AAV-GFP or AAV-GFP/Cre injections based on order of birth.

Immediately after the surgery, mice received subcutaneous analgesic (Carprofen 10 mg/kg). 24 hours after the surgery, mice received Metamizol in the drinking water (1.6 mg/ml). After surgery, mice were housed in pairs and were allowed to recover for 6 weeks before behavioral analysis (see below). Mice were sacrificed following behavioral analysis for histological verification of the virus injection site.

### OF behavior following virus injections

Open field behavior following stereotaxic virus injection was performed exactly as previously described (15, 18). Six weeks following surgery, mice were placed into a square open field chamber made of white plastic (50 x 50 cm) with a light intensity of 20 lux, and they were permitted to explore the chamber for 10 min. Performance was recorded using an overhead camera system and scored automatically using the Viewer software (Biobserve, St. Augustin, Germany). Additionally, the videos were analyzed in the Bonsai software to extract speed and location. We consider freezing behavior if the animal moved with a speed < 2 cm/s. Between each mouse, the arena was cleaned thoroughly with 70% ethanol followed by water to eliminate any odors left by the previous mouse.

### Statistical analysis

Data were analyzed statistically using MATLAB (*in vivo* electrophysiology) or GraphPad Prism (AAV-GFP/Cre injections). *In vivo* electrophysiological data were analyzed using non-parametric statistics. For comparison within groups, we used the Wilcoxon signed rank test and for comparison across groups we used two-sided Wilcoxon rank sum test. For analysis of the AAV-GFP/Cre data, outliers were first identified and excluded using the Grubb’s test (32), and normal distribution of the data was confirmed using the D’Agostino-Pearson normality test. Data were then analyzed using unpaired, two-tailed Student’s t-tests.

## Results

### Deletion of Nlgn2 alters LFP power throughout the anxiety network

To assess the consequences of Nlgn2 deletion on information processing throughout the anxiety network, we simultaneously recorded local field potential (LFP) oscillations in the mPFC, vHPC, BLA, CeM and BNST of male WT and Nlgn2 KO mice during exploration of an anxiogenic open field (OF) arena (Figure 1A and Figure S3). Under these conditions, Nlgn2 KO mice showed a pronounced anxiety phenotype as previously reported (15, 17, 18), including a reduction in time spent, distance traveled, and entries made into the center of the circular OF (Figure 1B-F). Spectral analysis of LFPs during OF exploration (including only periods of exploration with a speed of > 7 cm/s to eliminate potential locomotor influences) revealed distinctive oscillatory activity patterns in WT and Nlgn2 KO mice (Figure 1G). In WT mice, exposure to the OF resulted in a robust activation of the tripartite anxiety network consisting of the vHPC, the mPFC, and the BLA (23) (Figure 1H-J), with an increase in LFP power in the theta frequency range as previously reported (19, 22, 23) (Figure 1H), as well as a reduction in gamma power (Figure 1J). In contrast, this activation was less pronounced in the Nlgn2 KO mice, with no significant effect on theta power in mPFC or on gamma power in vHPC or BLA. Instead, we observed a significant activation of the nuclei that mediate defensive behaviors, i.e. the CeM and the BNST, with an increase in LFP power in the theta and beta frequency ranges (Figure 1H-J).

**Figure 1.**
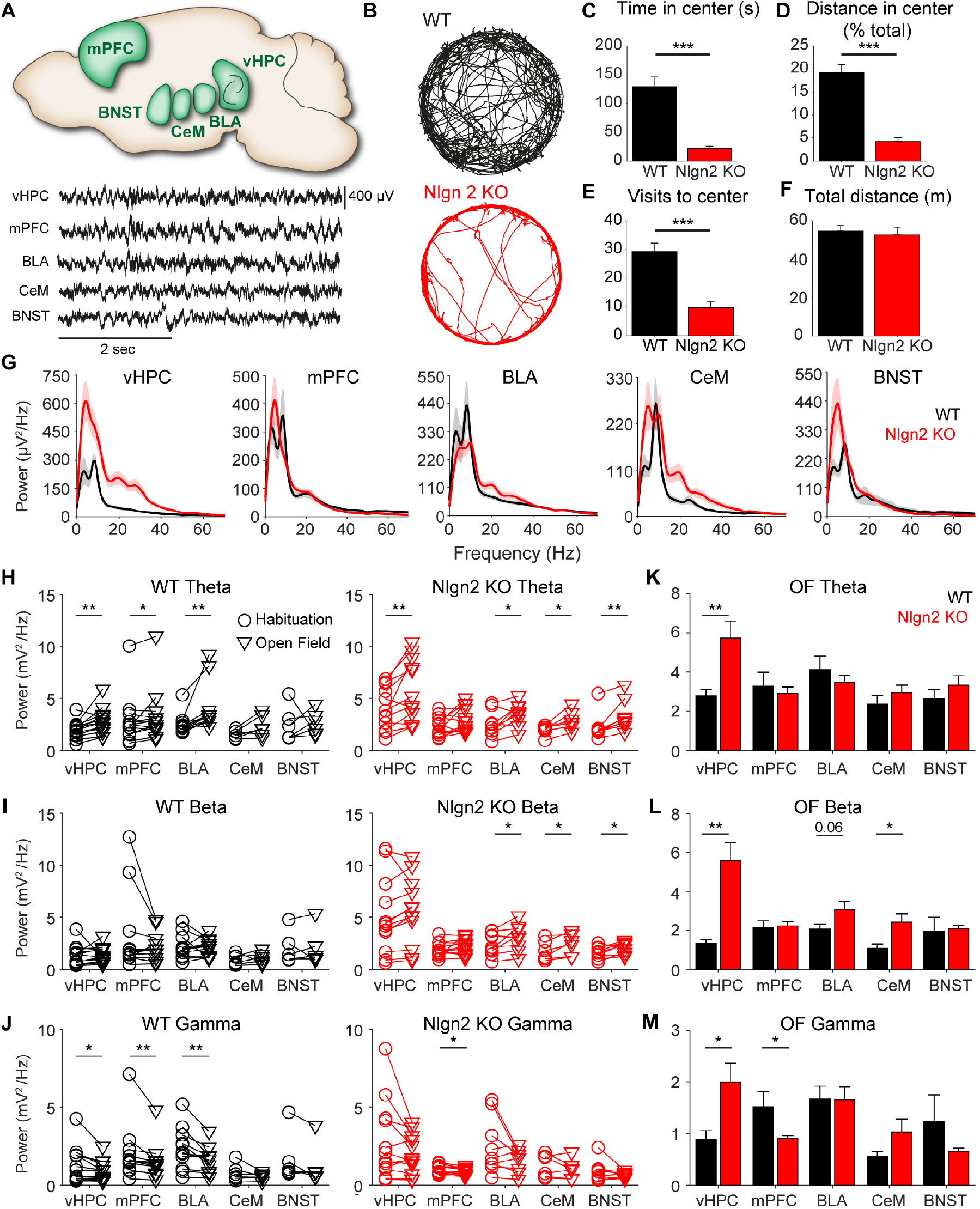
Deletion of Nlgn2 alters neuronal activity throughout the anxiety network. **(A)** Top, schematic of the brain regions recorded with a circuit approach (adapted from Calhoon & Tye (3)). Bottom, example of LFP traces recorded during exploration of the OF. **(B)** Representative tracks of the OF exploration. **(C)** Quantification of time spent in the anxiogenic center of the OF. **(D)** Distance travel in the center of the OF, expressed as a percentage of the total distance travelled. **(E)** Visits to the center of the OF. **(F)** Total distance travelled in the OF. **(G)** LFP power spectrum of the different brain regions recorded during OF exploration. **(H)** Theta power of WT (left) and Nlgn2 KO (right) animals during the habituation condition and OF exploration. **(I)** Beta power of WT (left) and Nlgn2 KO (right) animals during the habituation condition and OF exploration. **(J)** Gamma power of WT (left) and Nlgn2 KO (right) animals during the habituation condition and OF exploration. **(K)** Average theta power during OF exploration. **(L)** Average beta power during OF exploration. **(M)** Average gamma power during OF exploration. * p < 0.05, ** p < 0.01. n = 6-14 mice per genotype. Error bars represent SEM.

Comparison of LFP power in the OF in WT and Nlgn2 KO mice revealed a striking increase in power in the vHPC of Nlgn2 KO compared to WT mice that spanned all frequency ranges (Figure 1K-M). Moreover, more modest differences were observed in beta power in the CeM and the BLA (consistent with our previous report (18)) and in gamma power in the mPFC. Interestingly, the differences in power in the vHPC and mPFC were present even during the habituation phase before OF exposure (Figure S4A-C), indicating either that the Nlgn2 KO mice were already more anxious than WT mice during habituation (e.g. due to handling during attachment of the head stage) or that these differences were not immediately related to the anxiety phenotype. In contrast, the increased power observed in the BLA, CeM and BNST of Nlgn2 KO mice during OF exposure appeared to be specific to the OF context, since there were no significant genotype differences in these regions during habituation (Figure S4A-C). Together, these data indicate that deletion of Nlgn2 results in a robust increase in activity in the vHPC across multiple contexts as well as an abnormal recruitment of the BLA-CeM-BNST network during OF exploration.

### Deletion of Nlgn2 alters theta / beta frequency synchrony during OF exploration

Information processing in the brain relies critically on efficient communication between brain regions within a network, and indeed, disruptions in the functional connectivity between brain regions are thought to play a role in several psychiatric disorders (33–35), including anxiety disorders (36, 37). Such functional connectivity can be reflected in the synchronization of oscillatory activity across regions (33–35). To determine whether the anxiety phenotype observed in Nlgn2 KO mice was accompanied by alterations in neural synchrony, we measured coherence between brain regions both in the habituation context (Figure S4D-F) and during OF exploration (Figure 2A-E and Figure S5A-D) and calculated the average coherence for each brain region. Despite the fact that theta power was increased in Nlgn2 KO mice both in the transition from the habituation context to the OF and during the OF in comparison to WT mice (Figure 1H, K), coherence between brain regions in the theta frequency range was consistently lower in Nlgn2 KO mice (Figure 2B-C). However, this decrease in theta coherence was already present in the habituation context, particularly in vHPC, BLA and BNST (Figure S4D). Interestingly, we did not observe a modulation of theta coherence in transition from the habituation context to the OF in any of the genotypes (Figure S5B). In contrast, beta and gamma coherence was robustly decreased in WT mice in the OF compared to the habituation context (Figure S5C-D). Strikingly, this decrease in coherence was much less pronounced in Nlgn2 KO mice, with no significant decrease in beta coherence in any brain region and significant decreases in gamma only in the vHPC and mPFC. Consequently, beta coherence was significantly increased throughout the extended anxiety circuit during OF exploration in Nlgn2 KO mice compared to WT mice (Figure 2D). This increase in beta coherence appears to be specific to the anxiogenic OF, since no genotype differences were observed in the habituation condition (Figure S4E). As a consequence, the theta/beta frequency ratio, which has previously been linked to anxiety and attentional traits in human subjects (38), was increased from the habituation condition to the OF only in WT mice (Figure 2F-G) and Nlgn2 KO mice showed a significantly lower theta/beta ratio in the OF compared to WT mice (Figure 2H). These data indicate that altered functional connectivity between brain regions, and particularly the shift in coherence from the theta to the beta frequency range, may underlie the anxiety phenotype induced by deletion of Nlgn2.

**Figure 2.**
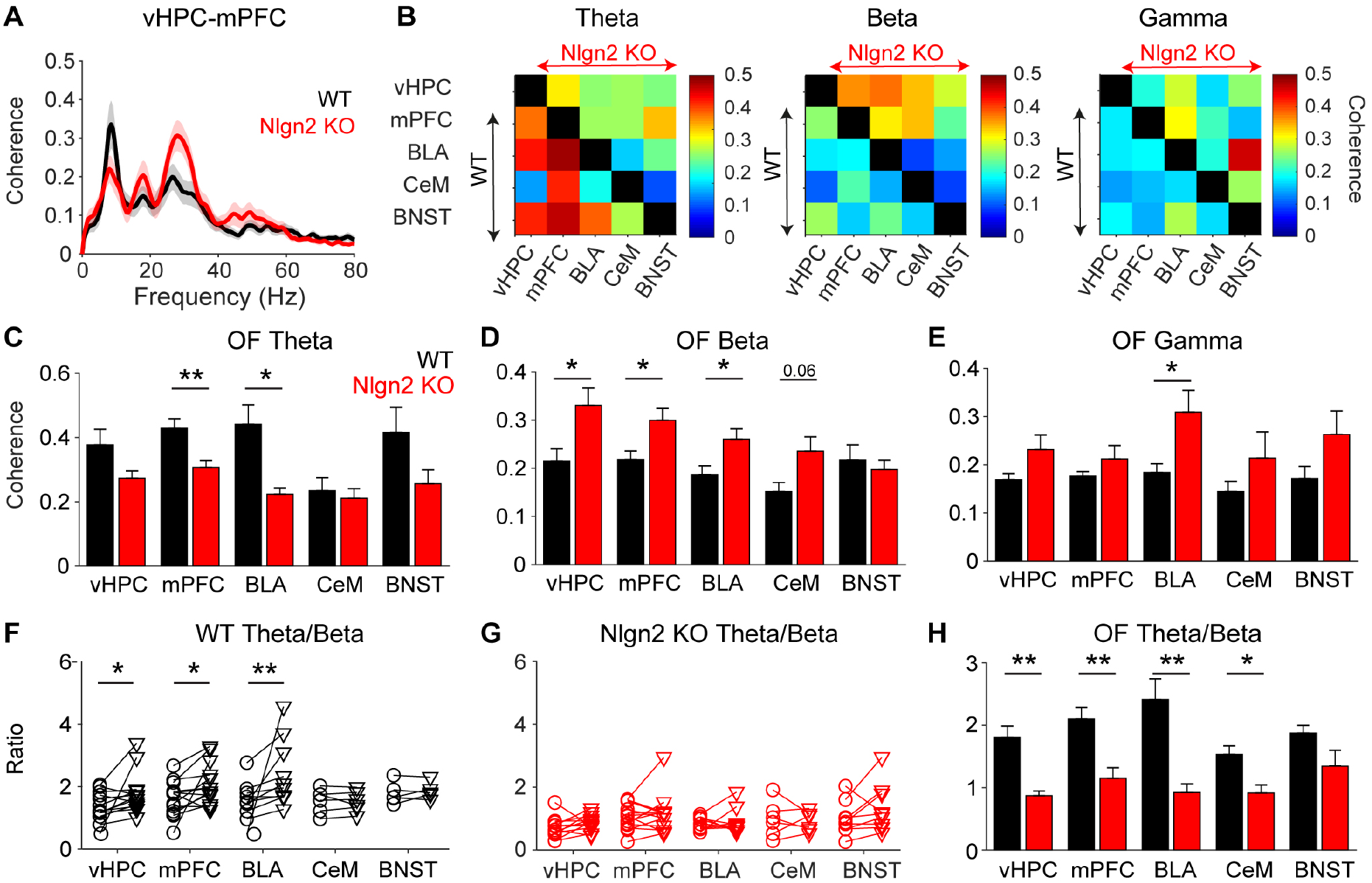
Deletion of Nlgn2 decreases theta/beta coherence in the anxiety circuit. **(A)** Example of average coherence for pairs of brain regions (here vHPC-mPFC, see Figure S5A for all pairs) simultaneously recorded during exploration of the open field. **(B)** Average coherence matrices between pairs of brain regions in the theta, beta and gamma frequency ranges in WT (lower left half of each matrix) and Nlgn2 KO (upper right half of each matrix) mice. **(C)** Average theta coherence during OF exploration. **(D)** Average beta coherence during OF exploration. **(E)** Average gamma power during OF exploration. **(F-G)** Theta/beta ratio for each brain region for WT (F) and Nlgn2 KO (G) animals during the habituation condition and OF exploration. **(H)** Average theta/beta ratio during OF exploration. * p < 0.05, ** p < 0.01. n = 6-13 mice per genotype. Error bars represent SEM.

### The theta / beta synchrony ratio best predicts anxiety behavior in the OF

To determine which of the above alterations may be most directly responsible for the anxiety phenotype in the Nlgn2 KO mice, we correlated anxiety behavior with LFP power and coherence between brain regions. Interestingly, we found that LFP power was a poor predictor of anxiety behavior, with only vHPC LFP power showing a significant correlation with entries into the center of the OF (Figure S6). In contrast, coherence in the beta frequency range, and in particular in the theta / beta frequency ratio, was strongly correlated with OF center entries throughout the anxiety network, with the most pronounced effects observed in the CeM and the BLA, followed by the mPFC and the vHPC (Figure 3C-D and Figure S7). These data support the notion that the shift in coherence from the theta to the beta frequency range may be the primary driving force behind the prominent anxiety phenotype in the Nlgn2 KO mice.

**Figure 3.**
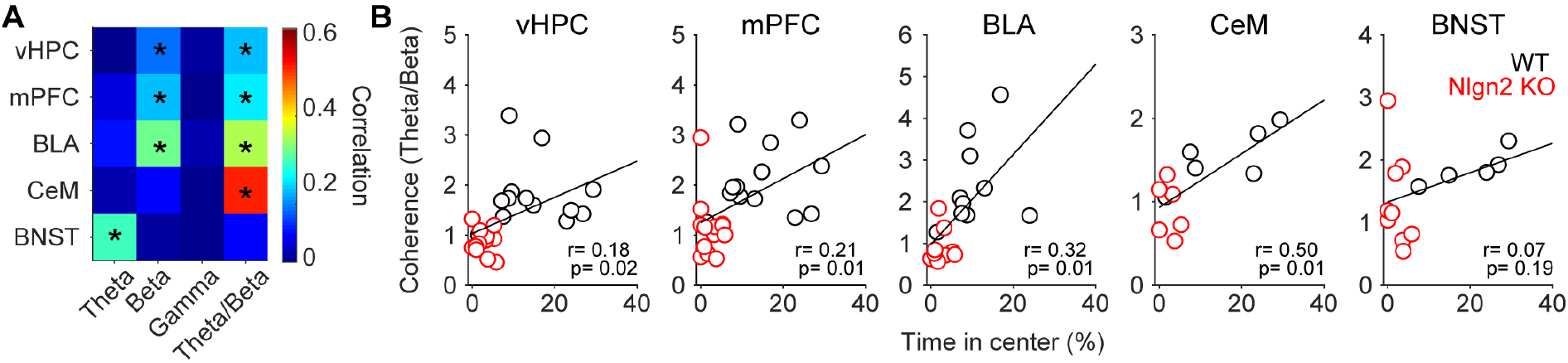
Theta/beta coherence strongly correlates with anxiety across brain regions. **(A)** Summary correlation matrix showing correlation between anxiety behavior and average coherence of each brain region in the theta, beta, gamma, and theta/beta frequency ranges. **(C)** Correlations of theta/beta coherence ratios with anxiety behavior for different brain regions. Statistical analysis of correlations is indicated in the figure. n = 6-13 mice per genotype.

### Deletion of Nlgn2 alters directed functional connectivity in the theta and beta frequency ranges during OF exploration

The above coherence analyses do not provide any information on the causality or directionality of the interactions between brain regions during anxiety processing. To specifically assess directionality in connectivity between vHPC, mPFC, BLA and CeM during OF exploration, we performed a Granger causality analysis as a measure of directed functional connectivity in the frequency domain (29, 30) (Figure 4A-J). In WT mice, high Granger causality values were observed in the vHPC→mPFC, vHPC→BLA and BLA→mPFC directions, specifically in the theta frequency range (Figure 4A-C, quantified in Figure 4K). These finding are consistent with previous reports indicating that the vHPC is causally involved in generating theta oscillations in the mPFC and BLA which contribute to the context-dependent regulation of anxiety-related behaviors (4, 19–23). In contrast, this leading role of vHPC theta oscillations was completely absent in Nlgn2 KO mice. Instead, increased Granger causality values were observed in the vHPC→BLA and mPFC→BLA directions in the beta frequency range (Figure 4B, H). Moreover, robust increases in beta directionality were observed in the vHPC→CeM, mPFC→CeM and CeM→vHPC directions (Figure 4D, E, F, I, quantified in Figure 4L), indicating that in the absence of Nlgn2, the vHPC and mPFC may exert its influence on the BLA and the CeM most strongly in the beta range. Together with the coherence measurements, these data suggest a model in which disinhibition in the vHPC of Nlgn2 KO mice leads to a loss of vHPC-directed theta synchrony and an increase in beta synchrony within the tripartite vHPC-mPFC-BLA network that may represent a potential circuit mechanism for the Nlgn2 KO anxiety phenotype (Figure 4M).

**Figure 4.**
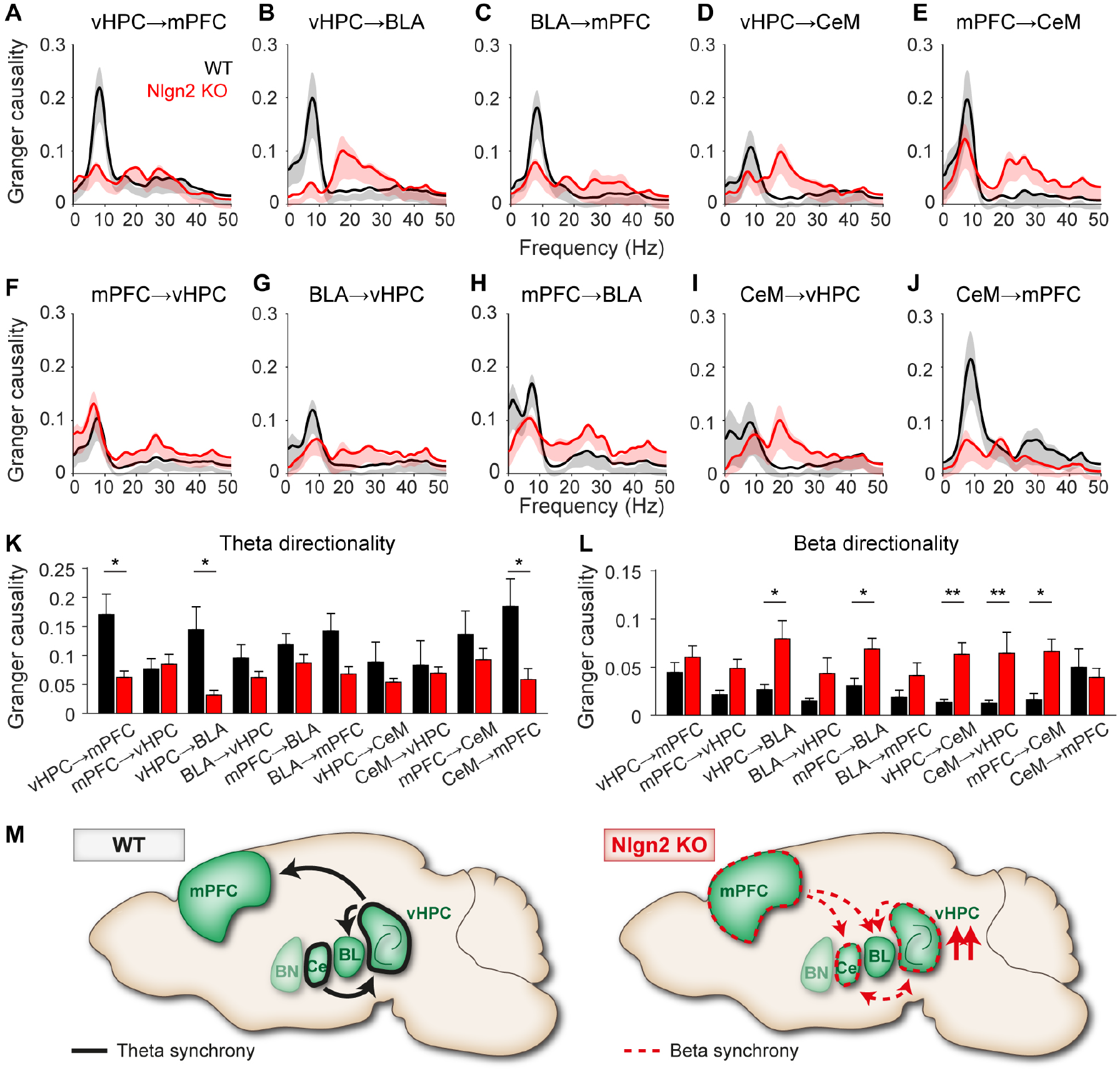
Nlgn2 deletion enhances vHPC-mPFC-CeM beta directionality. **(A-J)** Granger causality during OF exploration between (A) vHPC→mPFC, (B) vHPC→BLA, (C) BLA→mPFC, (D) vHPC→CeM, (E) mPFC→CeM, (F) mPFC→vHPC, (G) BLA→vHPC, (H) mPFC→BLA, (I) CeM→vHPC, (J) CeM→mPFC. **(K)** Average Granger causality in the theta range. **(L)** Average Granger causality in the beta range. * p < 0.05, ** p < 0.01. n = 5-12 mice per genotype. Error bars represent SEM. **(M)** Model of altered anxiety-related network activity in Nlgn2 KO mice. In WT mice, theta synchrony driven by the vHPC dominates anxiety-related network activity in the OF, consistent with previous reports (4, 19, 23). In Nlgn2 KO mice, disinhibition-induced increase of vHPC activity across frequency bands is accompanied by loss of vHPC-driven theta synchrony and increased beta synchrony distributed throughout the network. BL = basolateral amygdala, BN = bed nucleus of the stria terminalis, Ce = centromedial amygdala, mPFC = medial prefrontal cortex, vHPC = ventral hippocampus.

### vHPC and BLA encode distinct aspects of the Nlgn2 KO anxiety phenotype

To causally link the synaptic inhibition in each region to the anxiety phenotype, we generated Nlgn2 conditional KO (cKO) mice (see Figure S1 and S2 for details of generation and validation), and we subsequently deleted Nlgn2 locally in adult male homozygous cKO mice using stereotaxic injection of AAV vectors expressing Cre recombinase (AAV5-GFP/Cre) or a GFP control (AAV5-GFP). Given that it was previously already shown that local deletion of Nlgn2 in the mPFC is anxiolytic (39), likely reflecting the function of the mPFC in top-down control over the BLA, we focused our analysis on the vHPC (Figure 5A-M) and the BLA (Figure 5N-Z). To this end, we recorded OF behavior before and six weeks after the stereotaxic injection of AAV-Cre based on previous reports (39). We found that deletion of Nlgn2 from both the vHPC and the BLA resulted in an anxiety phenotype that partially resembled that of the constitutive Nlgn2 mice (Figure 1B-F and refs (15, 18)), but with different patterns. Specifically, deletion of Nlgn2 from the vHPC (Figure 5A-C) did not affect the behavior of the mice in the center of the OF (Figure 5D-H). Instead, it resulted in an increase in freezing behavior specifically upon transition from the safe periphery to the anxiogenic center, as reflected by an increase in the time spent in the transition zone (Figure 5J), a trend toward a reduction in the speed of movement in this zone (Figure 5K), and a pronounced increase in the time spent immobile (Figure 5M). In contrast, deletion of Nlgn2 from the BLA (Figure 5N-P) specifically resulted in a reduction in the number of visits to the center, likely reflecting anxiety-induced changes in exploratory behavior (15) (Figure 5R). These data indicate that the two brain regions both contribute to the alterations in anxiety processing resulting from Nlgn2 deletion but may encode separate aspects of the observed defensive behaviors, with the vHPC being critical for anxiety-induced freezing behaviors and the BLA for anxiety-induced reductions in exploration.

**Figure 5.**
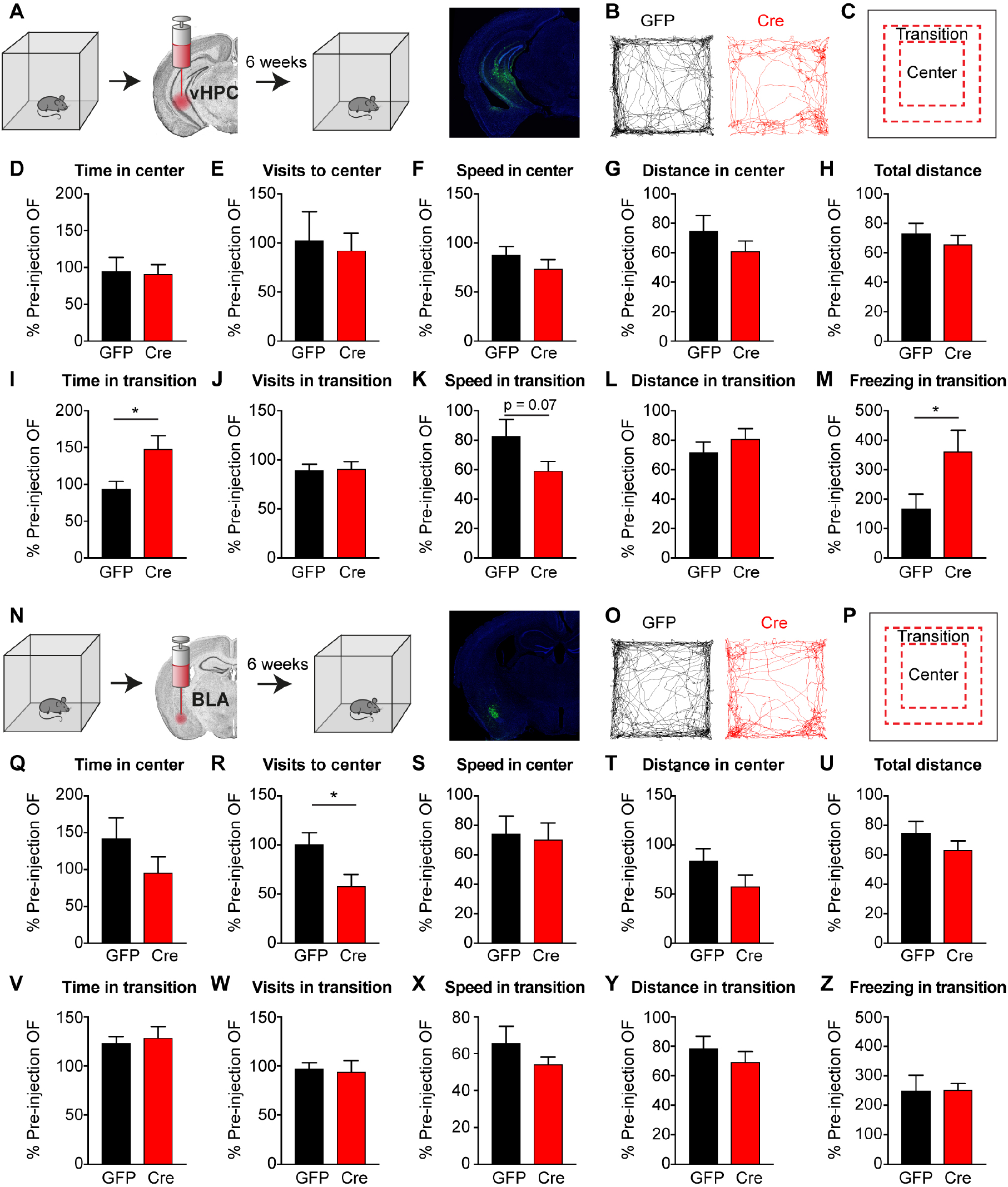
vHPC and BLA encode distinct aspects of the Nlgn2 KO anxiety phenotype. **(A)** Experimental design for stereotaxic injection of AAV-GFP or AAV-GFP/Cre into vHPC. Nlgn2 cKO mice were tested in the OF prior to and six weeks after stereotaxic injection. Correct targeting of virus was histologically confirmed following OF behavior. **(B)** Representative tracks of OF exploration. **(C)** Schematic representation of the center and transition zones analyzed. **(D)** Time in center of OF. **(E)** Visits to center of OF. **(F)** Speed in center of OF. **(G)** Distance in center of OF. **(H)** Total distance traveled in OF. **(I)** Time in transition zone. **(J)** Visits to transition zone. **(K)** Speed in transition zone. **(L)** Distance in transition zone. **(M)** Time freezing in transition zone. **(N)** Experimental design for stereotaxic injection of AAV-GFP or AAV-GFP/Cre into BLA. Nlgn2 cKO mice were tested in the OF prior to and six weeks after stereotaxic injection. Correct targeting of virus was histologically confirmed following OF behavior. **(O)** Representative tracks of OF exploration. **(P)** Schematic representation of the center and transition zones analyzed. **(Q)** Time in center of OF. **(R)** Visits to center of OF. **(S)** Speed in center of OF. **(T)** Distance in center of OF. **(U)** Total distance traveled in OF. **(V)** Time in transition zone. **(W)** Visits to transition zone. **(X)** Speed in transition zone. **(Y)** Distance in transition zone. **(Z)** Time freezing in transition zone. * p < 0.05. n = 6-12 per condition. Error bars represent SEM.

## Discussion

In the present study we show that excessive anxiety-related behavior in mice lacking in inhibitory synapse-associated protein Nlgn2 are encoded by alterations in functional connectivity between key brain regions in the anxiety network. In particular, pronounced increases in LFP power in the vHPC of Nlgn2 KO mice are accompanied by a loss of vHPC-directed theta frequency synchrony and an increase in beta frequency synchrony throughout the network. Moreover, altered theta / beta synchrony, but not LFP power in the individual brain regions, correlates strongly with anxiety behaviors. Local deletion of Nlgn2 from vHPC and BLA revealed distinct roles of Nlgn2 in these brain regions in encoding anxiety behaviors, with vHPC and BLA mediating freezing behavior and reduced exploratory activity, respectively. Together, our findings support a model in which impaired synaptic inhibition results in local disinhibition and profound alterations in the long-range functional connectivity underlying anxiety processing.

Perhaps the most striking observation arising from our study is that the exaggerated anxiety behavior in the Nlgn2 KO mice is most reliably predicted by the coherence in LFP oscillatory activity between brain regions, rather than the power of the oscillations in any given brain region. Nlgn2 KO mice displayed a dramatic increase in LFP power across frequency bands in the vHPC, as well as more modest increases in beta power in the CeM, consistent with our previous findings (18). No significant changes were observed in the BLA, although there was a trend for an increase in beta power, and in the mPFC, LFP power was reduced in the gamma frequency band. However, despite the fact that all of these brain regions have been prominently implicated in anxiety processing (1, 4, 5, 19–27, 40), only weak correlations were observed with time spent in the center of the OF as a measure of anxiety behavior (Figure S3). Instead, OF center time correlated most strongly with the synchrony between brain regions, and particularly with the ratio of theta/beta frequency synchrony, resulting from a reduction in theta frequency synchrony and a marked increase in synchrony in the beta frequency range. These findings indicate that the primary consequence of Nlgn2 deletion on information processing in the anxiety circuitry is to alter long-range communication between anxiety-related brain regions, thereby resulting in an aberrant activation of the CeM output to target regions that mediate defensive behaviors.

The observation that anxiety behaviors in Nlgn2 KO mice are accompanied by a reduction in theta frequency synchrony is surprising given that in WT mice, the exploration of an anxiogenic environment is accompanied by an increase in theta frequency synchrony (19, 20, 22, 23). It has been proposed that theta synchrony between the vHPC and the mPFC encodes certain aspects of the context relevant to anxiety, while theta synchrony between the mPFC and the BLA may contribute to the discrimination between safe and aversive contexts (4, 22, 23). Our data indicate that the prominent anxiety phenotype observed in the Nlgn2 KO mice does not result from an exaggerated activation of this normal vHPC-mPFC-BLA ‘tripartite anxiety network’ (23), but is instead accompanied by a shift towards abnormal coherence in the beta frequency range.

To date, relatively little is known about the role of beta oscillations in general, and to our knowledge, the role of beta frequency synchrony in anxiety processing has never been explored in animal models. Previous studies have identified a role for beta power in decision-making (41, 42) and context representation (43), as well as an increase in beta synchrony between the thalamus and the mPFC during working memory (44). We have recently shown that beta power is increased in the CeM of Nlgn2 KO mouse under anxiogenic conditions, and that this increase is normalized by a manipulation that also normalized the anxiety behavior (deletion of the adhesion protein IgSF9b) (18). Here, we expand these findings to indicate that beta synchrony throughout the vHPC-mPFC-BLA tripartite anxiety network may represent a neural signature of pathological anxiety processing in Nlgn2 KO mice. In support of this hypothesis, it was recently shown that amygdala-hippocampal beta coherence as well as the theta/beta ratio strongly correlates with anxiety traits in human subjects (38, 45).

The molecular basis for the observed alterations in LFP power or the theta/beta frequency synchrony shift remains to be elucidated. Nlgn2 is a key organizer protein at inhibitory synapses throughout the forebrain (8, 10, 13–16), and synaptic inhibition is well known to play a central role in the generation of local oscillatory activity (46–51). In particular, deletion of Nlgn2 is thought to most prominently affect perisomatic inhibitory synapses, likely those originating from parvalbumin (PV)-positive interneurons (10, 13–15) (although see also ref (52)). These synapses have been particularly implicated in the generation of oscillations in the gamma frequency range (47, 51, 53), and consistent with this notion, we previously showed that deletion of the Nlgn2-related Neuroligin-4 (Nlgn4) protein, which shares many properties with Nlgn2 at least in mice, results in pronounced alterations in gamma oscillatory activity in hippocampal slices (54). However, recent evidence indicates that PV-positive interneurons also contribute to the generation of theta and potentially beta frequency oscillations (46, 50, 55, 56), indicating that deletion of Nlgn2 may directly affect the power or coherence of these frequency ranges.

Whether these molecular alterations originate in one particular brain region, or whether they are distributed throughout the network, remains to be determined. Our data indicate that disinhibition in the vHPC may play a key role in the anxiogenic consequences of Nlgn2 deletion, resulting in dramatic increases in vHPC LFP power as well as in vHPC-directed functional connectivity during OF exploration. In accordance with this notion, local deletion of Nlgn2 from the vHPC resulted in a pronounced increase in freezing behaviors in the transition zone from the safe periphery to the anxiogenic center of the OF. Nevertheless, this manipulation does not recapitulate all of the anxiety-related consequences of constitutive Nlgn2 deletion, making it unlikely for the vHPC to be the sole origin of the anxiety phenotype (although other factors, such as limited virus spread or developmental effects in the constitutive KO, may also contribute to the differences). Indeed, local deletion of Nlgn2 in the BLA resulted in a significant reduction in visits to the center of the OF, consistent with the notion that Nlgn2 in the BLA may contribute specifically to the anxiety-induced reductions in exploratory behavior that we previously reported in Nlgn2 KO mice (15). Moreover, previous study showed that deletion of Nlgn2 in the mPFC resulted in an anxiety phenotype opposite to that observed in the constitutive Nlgn2 KO, with a reduction in avoidance behaviors in the OF and elevated plus maze (39), possibly reflecting the top-down regulatory role of the mPFC over the anxiogenic output of the amygdala. It is also possible that loss of Nlgn2 in brain regions not investigated here may contribute to the behavioral phenotype of the constitutive Nlgn2 KO mice. For example, local manipulations of Nlgn2 in the lateral septum (57) and nucleus accumbens (58) were recently shown to affect stress responsiveness and defensive behaviors.

Altogether, our findings indicate that deletion of Nlgn2 may have widespread consequences throughout the anxiety circuitry, and that the prominent anxiety-related behaviors induced by deletion of Nlgn2 are encoded by a fundamentally different mechanism than those in WT mice. Nlgn2 KO mice thus represent a highly relevant model to study the molecular and circuitry mechanisms by which alterations in synaptic inhibition may result in pathological anxiety behavior.

## Acknowledgments

The authors are grateful to Dr. Nils Brose for his continuous support of their research in the Department of Molecular Neurobiology, which is funded by the Deutsche Forschungsgesellschaft (CNMPB and SFB1190/P10) and the Bundesministerium für Bildung und Forschung (ERA-NET Neuron Synpathy). D.K.-B. was funded by a NARSAD Young Investigator Grant (Brain & Behavior Research foundation). O.B. was supported by a PhD fellowship from the Minerva Foundation. C.P.C. was a student of the Neurasmus Master program and was supported by an Erasmus Mundus scholarship (European Commission). V.M. was supported by an IMPRS stipend (Max Planck Society). N.B. was supported by a traineeship from the ERASMUS+ placement program (European Commission). The authors are grateful to Nikolas Karalis for sharing his code to calculate Granger causality and to Fritz Benseler, the AGCT Lab, the MPIEM animal facility and the Feinmechanik for excellent technical support.

## Author contributions

D.K.-B., H.C.-S. and O.B. conceived the study. H.C.-S. designed, performed and analyzed *in vivo* electrophysiology experiments. D.K.-B. and O.B. designed experiments involving local deletion of Nlgn2 in Nlgn2 cKO mice. O.B. and H.A. performed stereotaxic injection of AAV constructs into Nlgn2 cKO as well as subsequent behavioral experiments and data analysis. D.K.-B. and S.W. generated the Nlgn2 cKO mouse line. S.W., N.B. and H.A. performed molecular and behavioral validation of the Nlgn2 cKO mouse line. S.W. generated AAV particles for injection. C.P.C. assisted with stereotaxic surgeries and behavior experiments. V.M. assisted with *in vivo* electrophysiology and behavior experiments. D.K.-B. provided supervision for all experiments. D.K.-B., H.C.-S. and O.B. wrote the paper, and all authors edited and approved the final manuscript.

---

**Figure S1.**
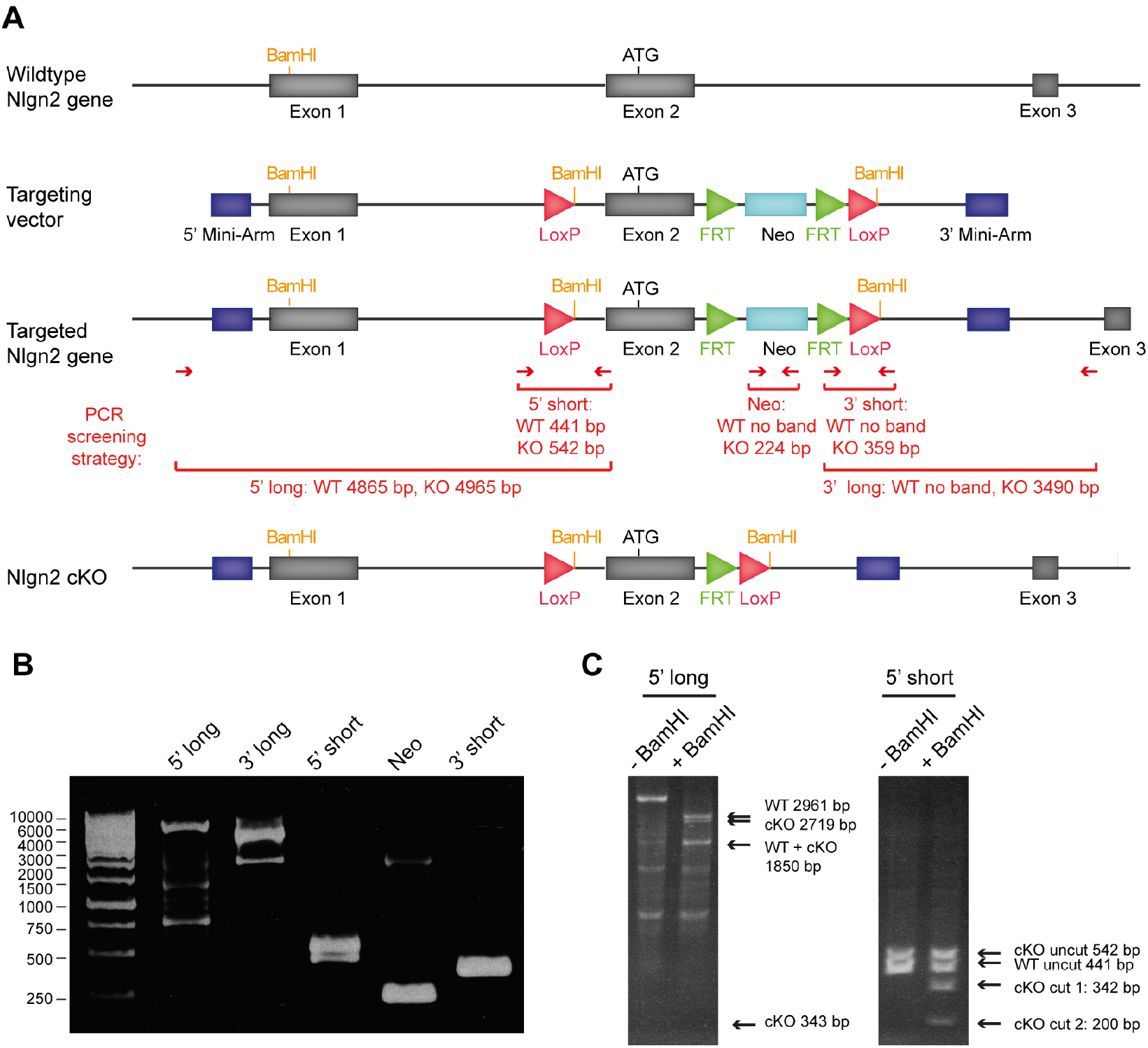
Generation of conditional Neuroligin 2 knockout mice. **(A)** Conditional Neuroligin-2 knockout mice (Nlgn2 cKO mice) were generated by insertion of loxP sites into the flanking regions of Nlgn2 exon 2 (which contains the start codon). LoxP sites were inserted using homologous recombination (59) based on the Nlgn2-containing BAC clone RPCIB731L16422Q(60) (ImaGenes GmbH). Targeted embryonic stem (ES) cells were generated using the wildtype C57BL/6N ES cell clone JM8A3 (61) (EUCOMM). Correct targeting of ES cell clones was confirmed by PCR analysis. Conditional KO mice were generated by blastocyst injection of targeted ES cell clone 8E02, and the Neo cassette was removed by crossing the mice with a Flp recombinase driver line. Nlgn2 cKO mice were backcrossed to a C57BL/6JRj background for at least six generations before experimental use. **(B)** PCR screening of ES cell clone 8E02. Five PCR products were generated using the primer sets depicted in panel a. All PCR bands corresponded to the bands expected from a correctly targeted ES cell clone. **c** BamHI digest of the PCR products “5’ long” and “5’ short” to confirm correct integration of the 5’ loxP site (which is associated with a BamHI site). Expected band sizes are marked with arrows. All products correspond to the bands expected from a correctly targeted ES cell clone.

**Figure S2.**
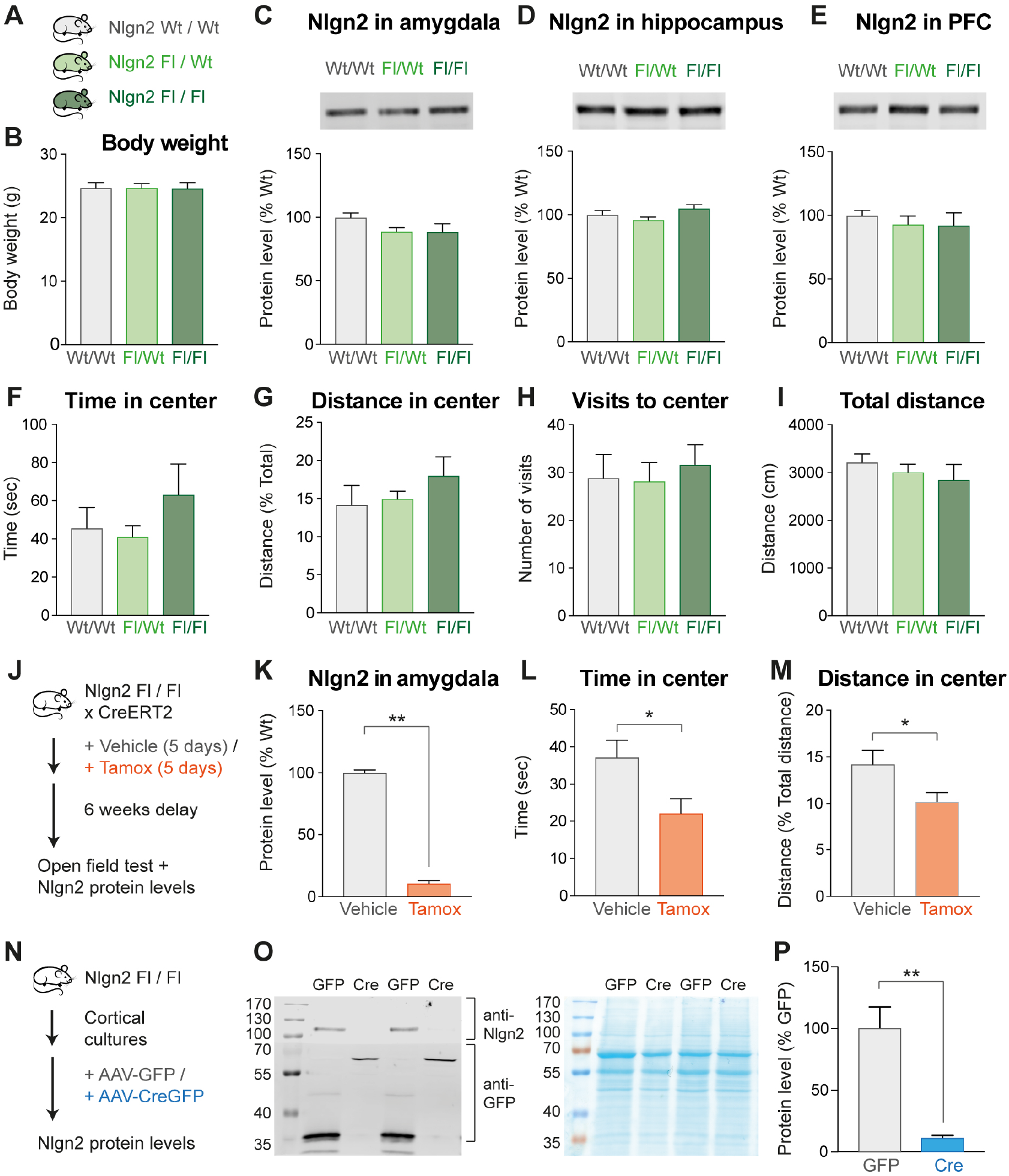
Validation of Nlgn2 cKO mice. **(A)** Nlgn2 cKO mice homozygous (Fl/Fl) or heterozygous (Fl/Wt) for the floxed allele were compared to WT littermates (Wt/Wt) in several measures relating to anxiety behavior. **(B)** No significant differences were observed between Wt/Wt, Fl/Wt and Fl/Fl mice in body weight. One-way ANOVA: F_2,21_ < 1. **(C-E)** Nlgn2 protein levels were determined in tissue homogenate from Wt/Wt, Fl/Wt and Fl/Fl mice as previously described (54, 62). No significant differences were observed in Nlgn2 protein levels in the amygdala (C, One-way ANOVA: F_2,15_ = 1.94, p = 0.178), hippocampus (D, One-way ANOVA: F_2,15_ = 2.81, p = 0.091) or prefrontal cortex (E, One-way ANOVA: F_2,14_ = 0.378, p = 0.692). **(F-I)** Anxiety behaviors were assessed in an open field test as previously described (15, 18). No significant differences were observed in time spent in the anxiogenic center (F), distance traveled in the center (G), visits to the center (H) or total distance traveled (I). One-way ANOVA: F_2,21_ < 1 for all behavioral comparisons. **(J)** To confirm efficacy of Nlgn2 deletion and emergence of the anxiety phenotype following deletion of Nlgn2 in adulthood, Nlgn2 cKO (Fl/Fl) mice were crossed with a Tamoxifen-inducible Cre driver (ROSA26_CreERT2) generated in the laboratory of Tyler Jacks (63) and obtained from the Jackson Laboratory). Tamoxifen dissolved in corn oil was injected intraperitoneally for 5 consecutive days, 2x per day at 100 mg/kg. Six weeks after administration of Tamoxifen, mice were tested in the open field and subsequently sacrificed for analysis of Nlgn2 protein levels. **(K)** Nlgn2 protein levels in the amygdala were strongly reduced following Tamoxifen administration. **(L-M)** Following Tamoxifen administration, Nlgn2 cKO spent significantly less time (L) and traveled less distance (M) in the center of the open field, confirming that deletion of Nlgn2 in adulthood has an anxiogenic effect. **(N-P)** To confirm efficacy of Nlgn2 deletion by AAV-Cre viruses, dissociated cortical cultures were infected with AAV-GFP or AAV-CreGFP 24 hour after plating. After 14 days, cultures were harvested and Nlgn2 protein levels were determined by immunoblotting. Nlgn2 levels were significantly reduced following infection with AAV-CreGFP compared to infection with AAV-GFP, confirming the AAV-CreGFP viruses were effective in deleting Nlgn2. All bars represent mean + SEM, * p < 0.05, ** p < 0.01, *** p < 0.001. n = 6-8 for Nlgn2 cKO experiment, n = 10-11 for Tamoxifen experiment, n = 4 for AAV-CreGFP experiment.

**Figure S3.**
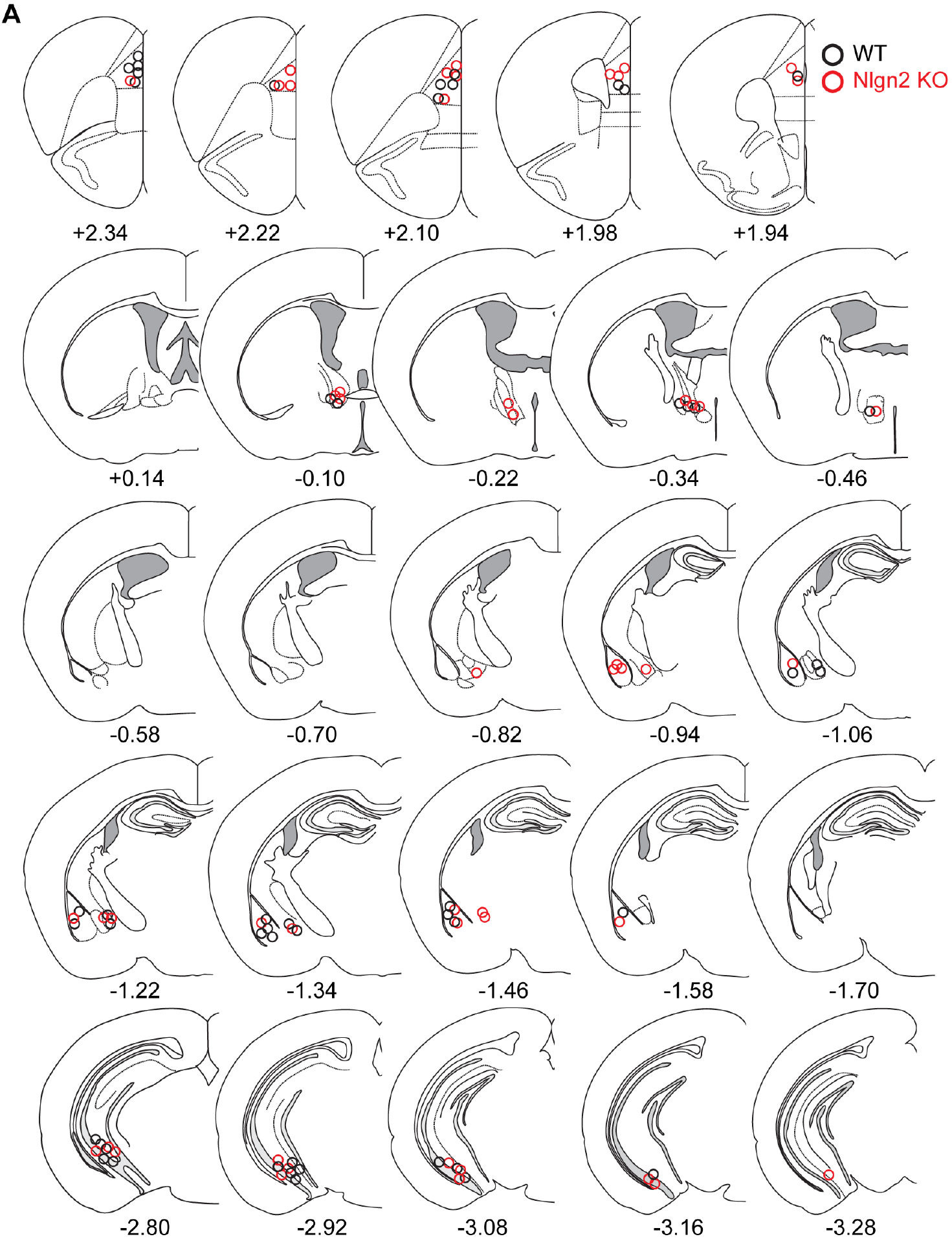
Histological verification of *in vivo* electrophysiology recording sites. **(A)** Black and red circles represent location of the electrode tips in WT and Nlgn2 KO mice, respectively.

**Figure S4.**
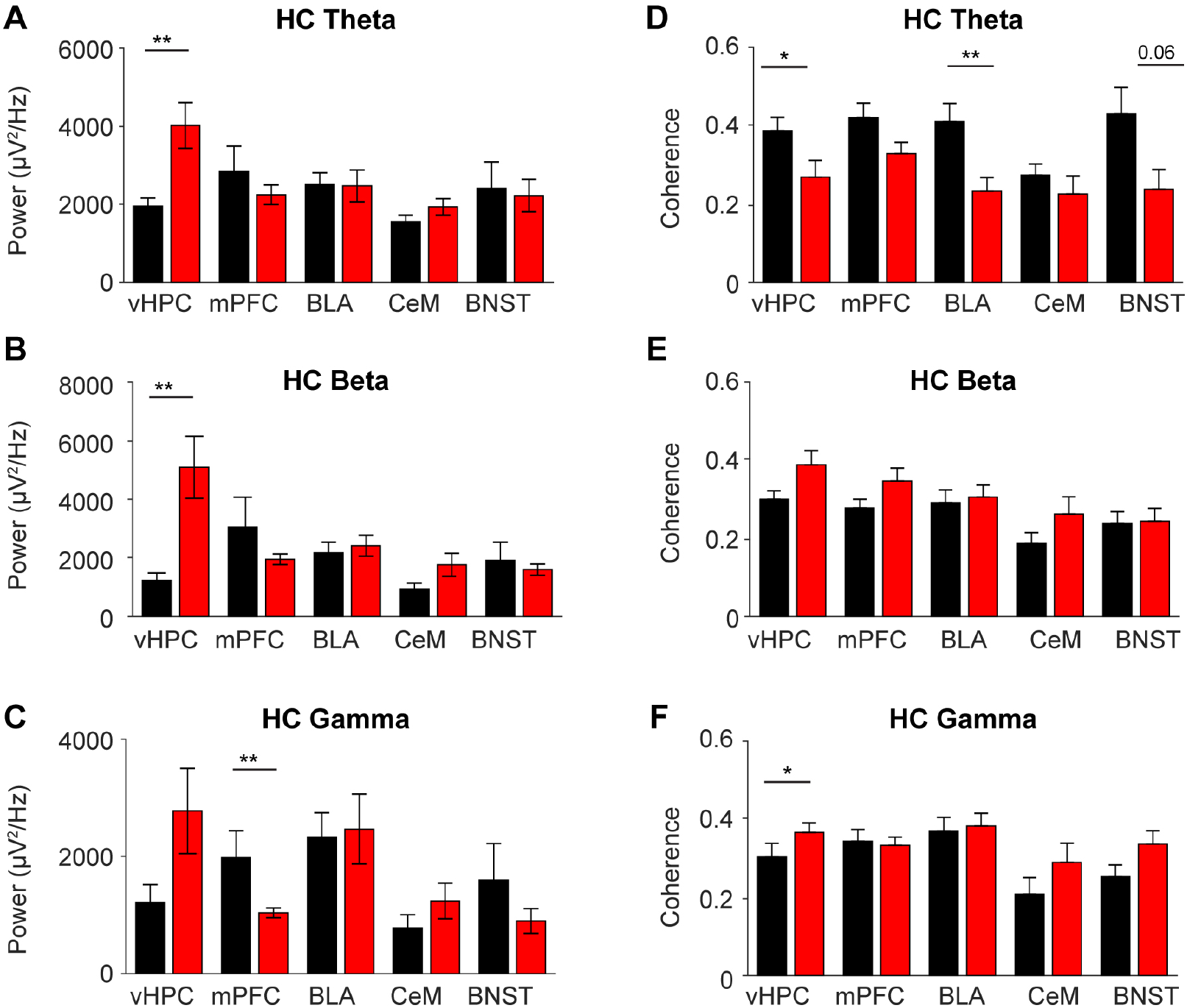
LFP power and coherence in WT and Nlgn2 KO mice in the habituation context. Mice were tethered and recorded in the habituation context (home cage with dim light, ~ 30 lux) for 5 min. **(A-C)** Average power during habituation in the theta (A), beta (B) and gamma (C) frequency ranges. **(D-F)** Average coherence during habituation in the theta (D), beta (E) and gamma (F) frequency ranges. * p < 0.05, ** p < 0.01. n = 6-14 mice per genotype. Error bars represent SEM.

**Figure S5.**
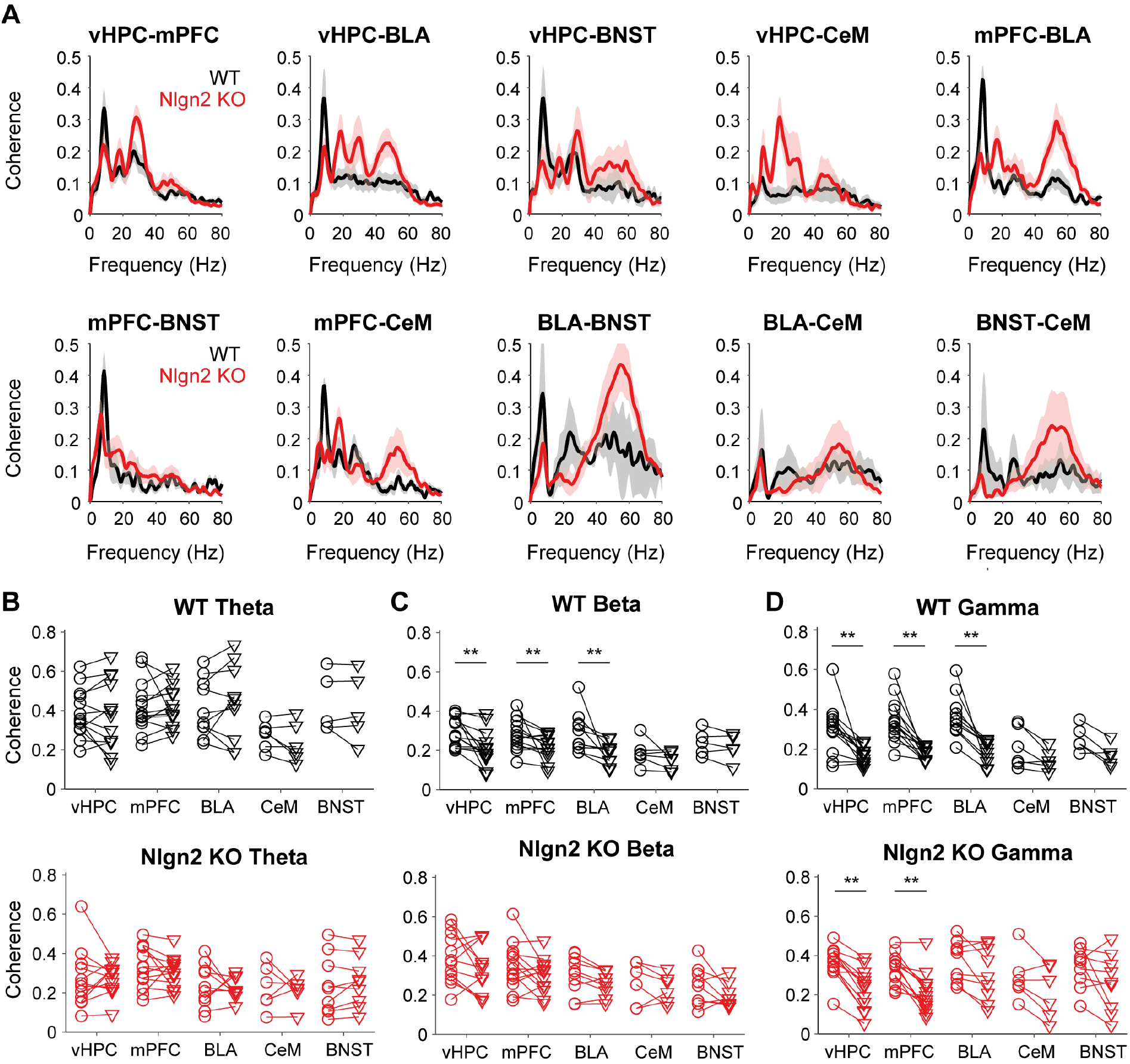
Coherence during exploration of the OF in WT and Nlgn2 KO mice. **(A)** Average coherence for pairs of brain regions simultaneously recorded during exploration of the open field. **(B-D)** Average coherence in the theta (B), beta (C) and gamma (D) frequency range for each brain region for WT (upper panel) and Nlgn2 KO (lower panel) animals during the habituation condition and OF exploration. ** p < 0.01. n = 6-13 mice per genotype. Error bars represent SEM.

**Figure S6.**
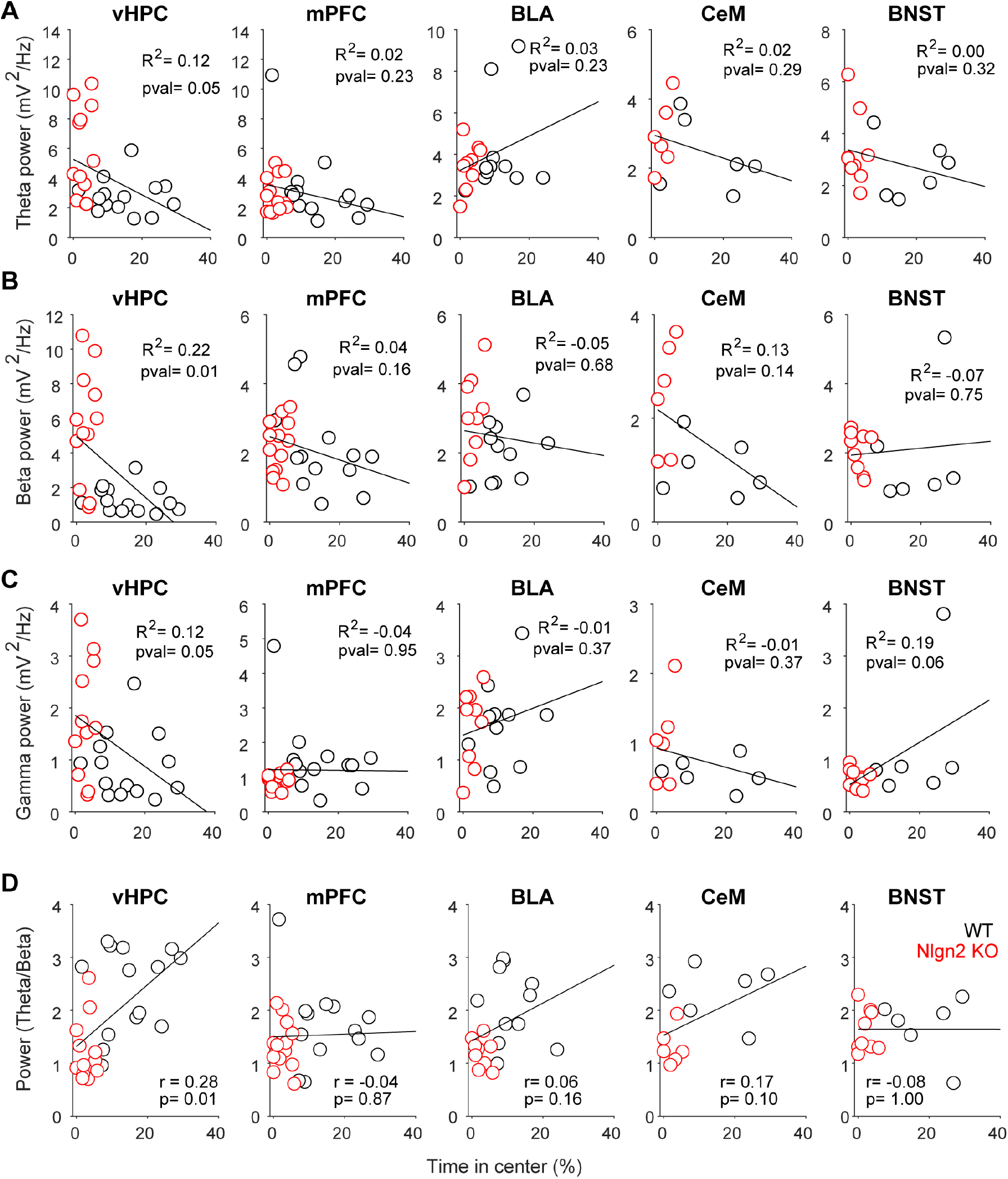
Correlation between LFP power and anxiety behavior during exploration of the OF. **(A-D)** Correlation between anxiety behavior and average power in the theta (A), beta (B), gamma (C) and theta/beta (D) frequency ranges. Statistical analysis of correlations is indicated in the figure. n = 6-14 mice per genotype.

**Figure S7.**
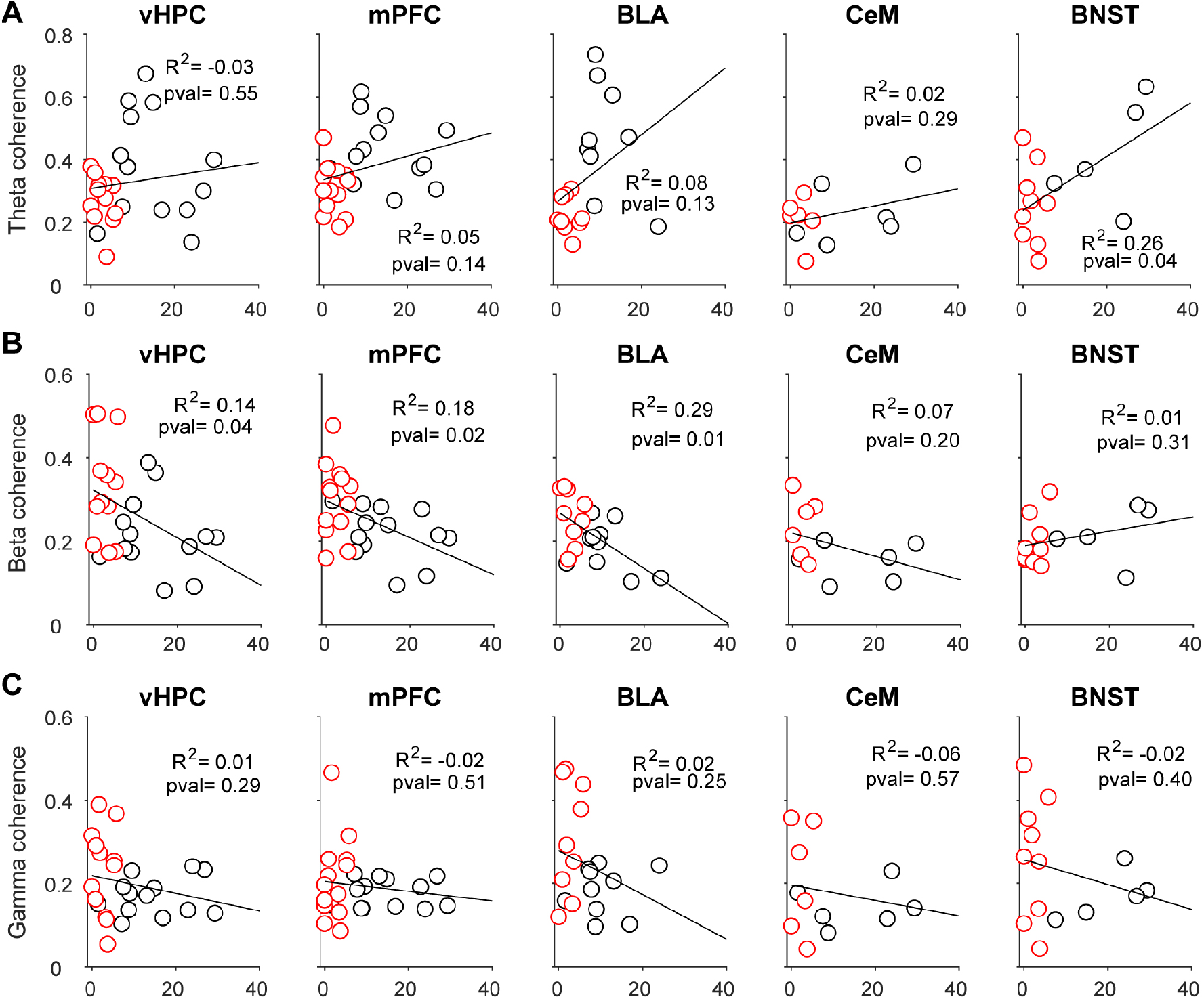
Correlation between LFP coherence and anxiety behavior during exploration of the OF. **(A-C)** Correlation between anxiety behavior and average coherence in the theta (A), beta (B) and gamma (C) frequency ranges. Statistical analysis of correlations is indicated in the figure. n = 6-14 mice per genotype.

